## Supplementary for "Fast Viral Dynamics Revealed by Microsecond Time-Resolved Cryo-EM"

### Materials and Methods

Sample preparation

Cowpea chlorotic mottle virus was provided by the group of Prof. Jeroen Cornelissen at the University of Twente. Cryo samples of contracted CCMV (pH 5, free of divalent ions were prepared by applying 3 µl of the sample solution (11.5 mg/ml in 100 mM sodium acetate buffer, 1 mM sodium azide, 1 mM EDTA) to glow discharged UltrAuFoil grids (R1.2/1.3, 300 mesh, Quantifoil), which were plunge frozen with a Thermo Fisher Vitrobot Mark IV (10 °C, 95 % relative humidity, 3 s blotting time, blotting force of 10). Cryo samples of extended CCMV (pH 7.6) were prepared from the sample solution through buffer exchange with a centrifugal concentrator (MWCO 50 kDa, 10°C). The sample solution (23 mg/ml in 10 mM Tris buffer, 50 mM sodium chloride, 1mM EDTA) was then plunge frozen under the same conditions. For pH jump experiments, NPE-caged-proton (Bio-Techne, 120 mM) was added to this solution in a 1:6 volume ratio prior to plunge freezing.

In order to release the photoacid, cryo samples were placed in a Linkam cryo stage and irradiated with a nanosecond pulsed UV laser for 30 min (Bright Solutions Wedge, 266 nm, 35 mW, 7.1±0.1 mm FWHM spot size in the sample plane, as determined from a knife edge scan). The pH of the sample after release of the photoacid was determined as follows. A frozen droplet of the same sample solution (3 µl) was UV irradiated under identical conditions, after which the droplet was melted. A measurement with a pH strip yielded a pH of 4.5± 0.5.

In order to determine that UV irradiation does not cause contraction of the particles in the absence of a photoacid (Fig. S2), cryo samples of extended CCMV (without photoacid) were irradiated under identical conditions.

In-situ revitrification

Revitrification experiments were performed *in situ* with a modified JEOL 2200FS transmission electron microscope (*28*) as previously described (*17*). Microsecond laser pulses for revitrification (532 nm wavelength, about 100 mW power) were obtained by chopping the output of a continuous laser (Laser

Quantum, Ventus 532) with an acousto-optic modulator (AA Opto-Electronic). The laser beam was focused to a spot size of 28±2 μm FWHM in the sample plane, as determined from a knife edge scan. Sample areas were melted and revitrified with the laser beam centered onto a grid square.

Data collection

Data were collected at the Dubochet Center for Imaging in Lausanne using a Titan Krios microscope equipped with Falcon IV detector and a SelectrisX energy filter. The data acquisition parameters for the different data sets are listed in Tables S1-3.

Single-particle reconstructions

Single-particle reconstructions were performed in CryoSPARC 4.0.1 (*20*). The following workflow was used, with details for the different data sets are provided in Figs. S1 and S2. The micrographs were patch motion corrected, and movies with a total full-frame motion distance of more than 100 pixels were discarded. Contrast transfer function (CTF) estimation was performed using Patch CTF. Micrographs with an estimated resolution of less than 10 Å or an astigmatism of over 2000 Å were discarded, as well as micrographs containing hexagonal or cubic ice. Particles were picked using manual picking or blob picking, followed by template picking and were extracted with a box size of 768 pixels. Following 2D classification and *ab initio* reconstruction, the best classes resulting from heterogeneous refinement (I symmetry) were further refined using homogeneous or non-uniform refinement with I symmetry imposed. Additional refinement steps included CTF refinement, per particle defocus refinement, and Ewald sphere correction. The global and local resolution was estimated using an gold-standard FSC threshold of 0.143. The structures were visualized with UCSF ChimeraX 1.5 (*29*, *30*).

Variability analysis

The variability analysis shown in Fig. 3E was performed on a combined dataset containing 21,675 refined particles each of the extended, partially contracted, and fully contracted CCMV. Three variability components and a low-pass filter resolution of 10 Å were used. In Fig. 3E, the particle distribution is shown as a function of the first two components. To analyze the motions of the capsid proteins during contraction, the particle distribution was divided into 30 equal slices along the first component, and each slice was homogeneously reconstructed with icosahedral symmetry imposed.

Model building

A model of contracted CCMV capsid was obtained with the following procedure. The model of contracted CCMV from PDB:1ZA7 (*31*) was protonated using the APBS-PDB2PQR software suite (*32*) with the pH set to 5 and rigidly fit into the auto-sharpened contracted CCMV map using UCSF ChimeraX 1.4 (*29*, *30*). The asymmetric unit was then iteratively refined using the real space refine tool in PHENIX 1.20.1-4487 (*33*), with secondary structure and non-crystallographic symmetry constraints imposed, and the ISOLDE plugin (*34*) in UCSF ChimeraX. The refinement was finalized using a map section containing the asymmetric unit as well as the chains in its proximity.

A model of the extended CCMV capsid was obtained with a similar procedure. The model of the contracted form obtained above was set to the protonation state at pH 7.6 and rigidly fit into the auto-sharpened map of extended CCMV. The asymmetric unit was then refined as above while imposing torsional and adaptive distance restraints derived from the contracted structure. Imposing these restraints helped prevent unphysical geometries from occurring during the refinement in the 3.9 Å map. The quality of the models were evaluated using MolProbity (*35*) and PHENIX (*33*). The atomic models of the asymmetric units are shown in Fig S4.

Analysis of the motions of the capsid proteins during contraction

In order to analyze the motions of the capsid proteins during contraction, the particle distribution in Fig. 3E was divided into 30 slices along the first component, and a reconstruction was obtained for each slice. The model of the extended state was then rigidly fit into the reconstructions of slices 1–12 (extended and partially contracted configurations), and the model of the contracted state into the reconstructions of slices 25–30 (contracted configurations). The positions of the alpha carbons were then used to extract the geometric parameters reported in Fig. 4.

The capsid diameter (Fig. 4B) was measured along 5-fold symmetry axis and corresponds to the largest distance of two alpha carbons along this direction. The angles of the rotation of the pentamers around the 5-fold axis (Fig. 4C) were determined from the displacements of the centers of mass of the B proteins. The rotation angles of the hexamers around the 3-fold axis (Fig. 4C) represent an average of the rotation angles of the A and C proteins, which were calculated in the same manner. After subtracting the rotation of the capsomers, the superimposed rotation of each capsid protein around its center for mass was then calculated (Fig. 4D).

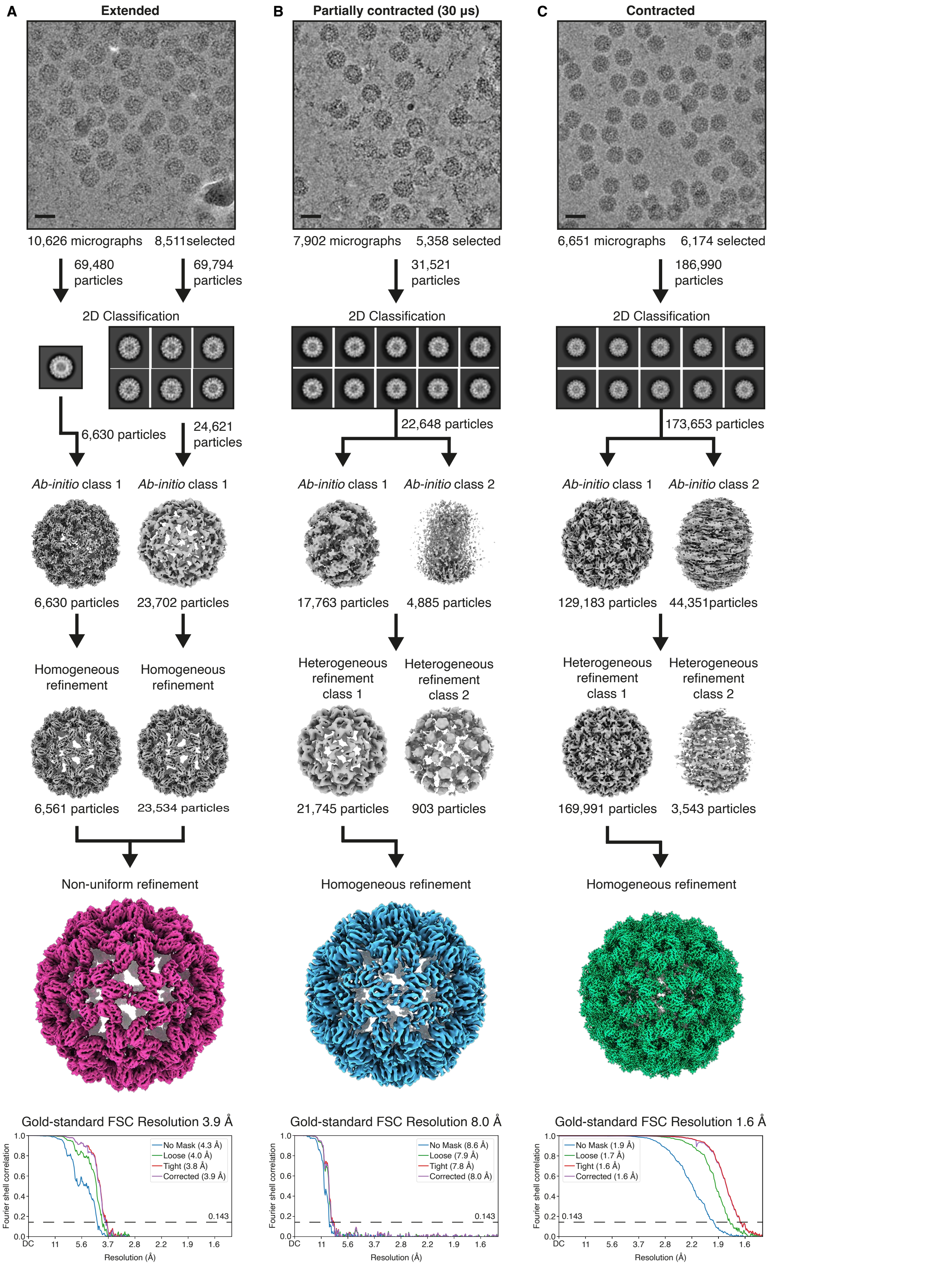
A
Extended
B
Partially contracted (30 μs)
C
Contracted
10,626 micrographs 8,511selected
7,902 micrographs 5,358 selected
6,651 micrographs 6,174 selected
69,480 particles
69,794 particles
31,521 particles
186,990 particles
2D Classification
2D Classification
2D Classification
6,630 particles
24,621 particles
22,648 particles
173,653 particles
Ab-initio class 1
Ab-initio class 1
Ab-initio class 1
Ab-initio class 2
Ab-initio class 1
Ab-initio class 2
6,630 particles
23,702 particles
17,763 particles
4,885 particles
129,183 particles
44,351particles
Homogeneous refinement
Homogeneous refinement
Heterogeneous refinement class 1
Heterogeneous refinement class 2
Heterogeneous refinement class 1
Heterogeneous refinement class 2
6,561 particles
23,534 particles
21,745 particles
903 particles
169,991 particles
3,543 particles
Non-uniform refinement
Homogeneous refinement
Homogeneous refinement
Gold-standard FSC Resolution 3.9 Å
Gold-standard FSC Resolution 8.0 Å
Gold-standard FSC Resolution 1.6 Å
No Mask (4.3 Å)
Loose (4.0 Å)
Tight (3.8 Å)
Corrected (3.9 Å)
No Mask (8.6 Å)
Loose (7.9 Å)
Tight (7.8 Å)
Corrected (8.0 Å)
No Mask (1.9 Å)
Loose (1.7 Å)
Tight (1.6 Å)
Corrected (1.6 Å)
0.143
Fourier shell correlation
Resolution (Å)
DC 11 5.6 3.7 2.8 2.2 1.9 1.6

**Fig. S1. Single-particle reconstruction workflows for the extended (A), partially contracted (B), and fully contracted configurations of the CCMV capsid (C).** Gold Standard Fourier Shell Correlations are shown for each data set, with the dashed black line indicating the 0.143 cutoff. For the extended configuration in (**A**), two datasets were processed separately and merged in the final reconstruction.

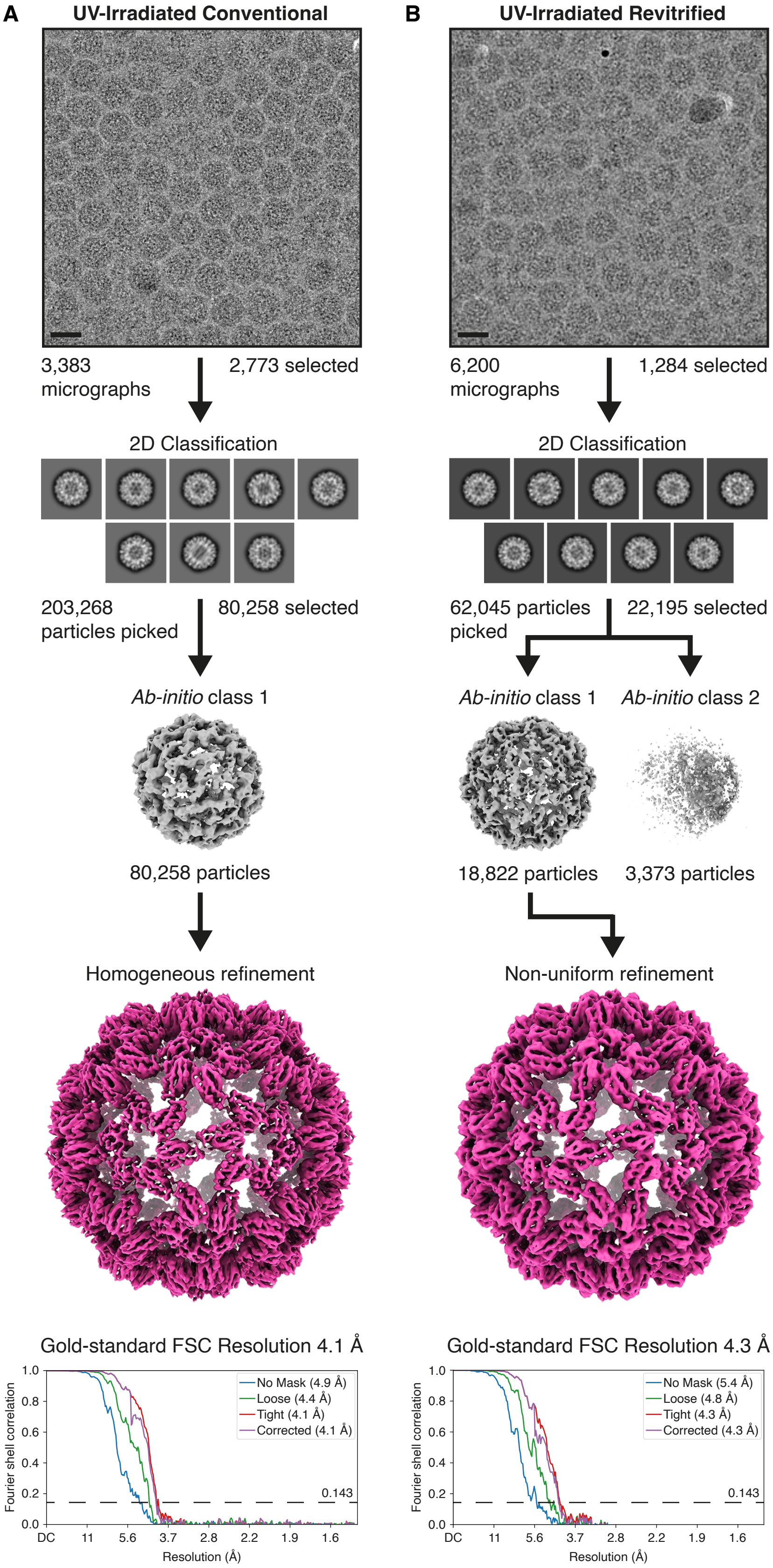
A
UV-Irradiated Conventional
B
UV-Irradiated Revitrified
3,383 micrographs
2,773 selected
6,200 micrographs
1,284 selected
2D Classification
2D Classification
203,268 particles picked
80,258 selected
62,045 particles picked
22,195 selected
Ab-initio class 1
Ab-initio class 1
Ab-initio class 2
80,258 particles
18,822 particles
3,373 particles
Homogeneous refinement
Non-uniform refinement
Gold-standard FSC Resolution 4.1 Å
Gold-standard FSC Resolution 4.3 Å
No Mask (4.9 Å)
Loose (4.4 Å)
Tight (4.1 Å)
Corrected (4.1 Å)
No Mask (5.4 Å)
Loose (4.8 Å)
Tight (4.3 Å)
Corrected (4.3 Å)
0.143
Fourier shell correlation
Resolution (Å)
DC 11 5.6 3.7 2.8 2.2 1.9 1.6

**Fig. S2. Irradiation with UV light in the absence of a photoacid does not cause CCMV to contract. Single-particle reconstruction workflows for a UV irradiated sample of the extended configuration of CCMV.** The sample was prepared at pH 7.5 in the absence of a photoacid and irradiated with UV light. No contraction is observed after revitrification. Gold Standard Fourier Shell Correlations are shown for each data set, with the dashed black line indicating the 0.143 cutoff.

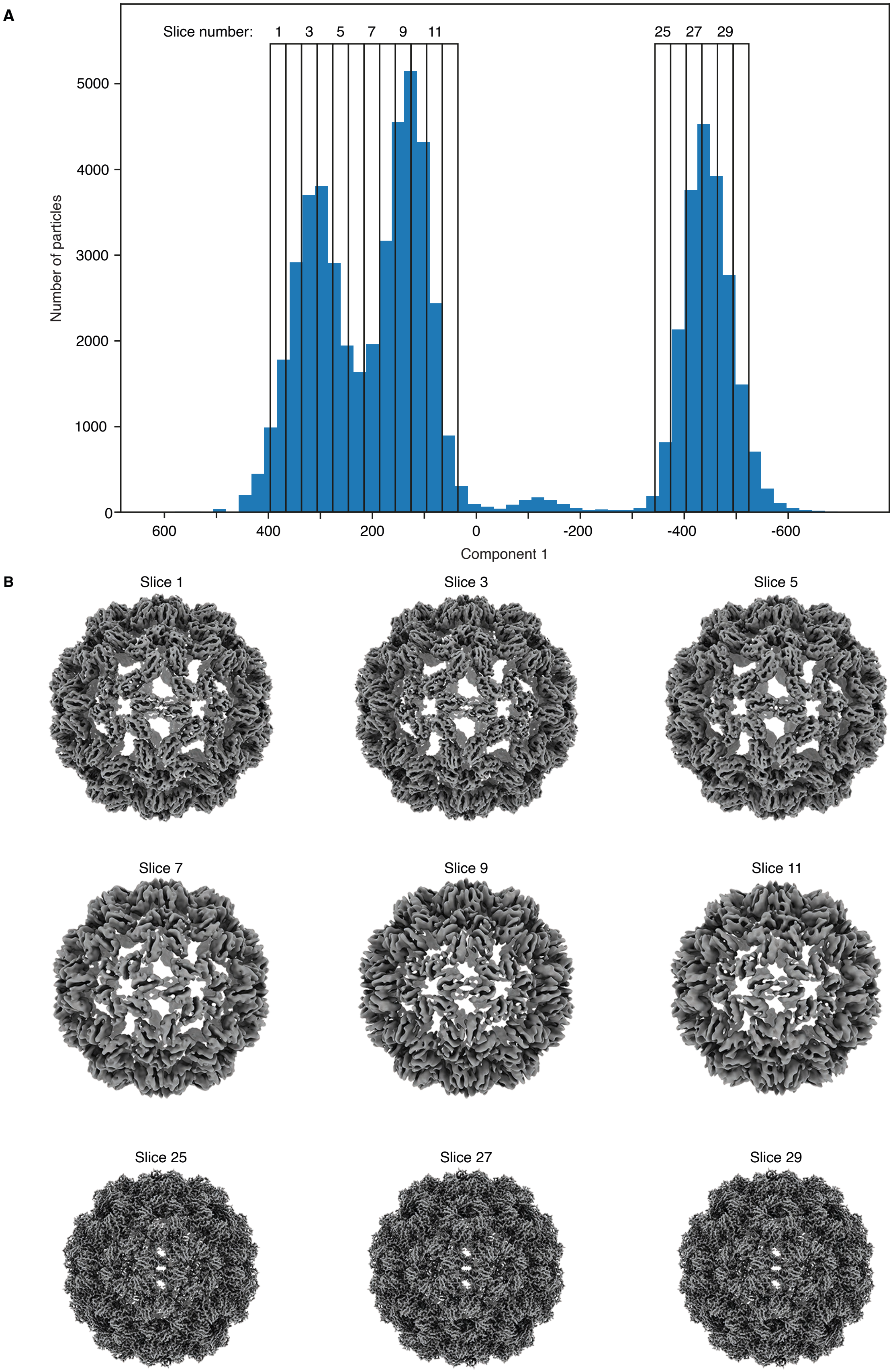
A
Slice number: 1 3 5 7 9 11 25 27 29
Number of particles
0
1000
2000
3000
4000
5000
600 400 200 0 -200 -400 -600
Component 1
B
Slice 1
Slice 3
Slice 5
Slice 7
Slice 9
Slice 11
Slice 25
Slice 27
Slice 29

**Fig. S3. Variability analysis of a combined dataset of the extended, partially contracted, and fully contracted configurations of CCMV from Fig. 3E. (A)** Particle distribution as a function of the first component. The distribution is divided into 30 slices along the first component. (**B**) Reconstructions from representative slices.

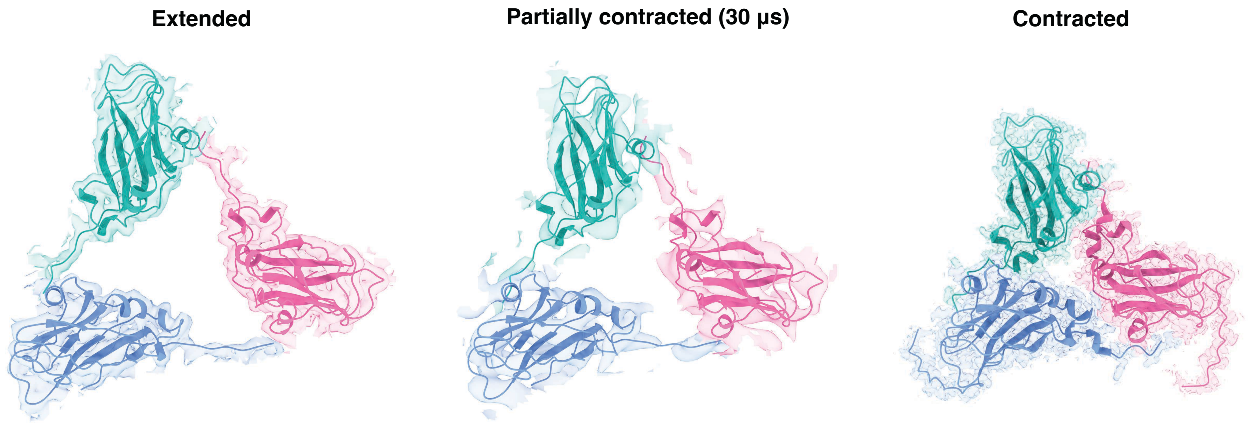


**Fig. S4. Models of the asymmetric unit of the extended, partially contracted, and fully contracted states.**

**Table S1: Contracted CCMV data collection parameters, and reconstruction and modelling statistics.**

| | |
|---|---|
| **Data collection and processing** | |
| Microscope | Thermo Fisher Titan Krios |
| Detector | Falcon 4 |
| Energy Filter | SelectrisX, 10 eV band width |
| Magnification | 165,000 |
| Accelerating voltage (kV) | 300 |
| Total electron exposure ($e^-/Å^2$) | 50 |
| Defocus (µm) | -0.3 – -0.9 |
| Defocus increment (µm) | 0.1 |
| Pixel size (Å) | 0.726 |
| Initial movies (no.) | 6,651 |
| Symmetry imposed | I |
| Initial particle images (no.) | 186,990 |
| Final particle images (no.) | 169,835 |
| Map resolution (Å) | 1.6 |
| FSC threshold | 0.143 |
| **Refinement** | |
| Map sharpening B factor ($Å^2$) | 40.7 |
| **Model composition** | |
| Non-hydrogen atoms | 3,593 |
| Protein residues | 479 |
| Ligands | n/a |
| **Validation** | |
| All-atom clashscore | 0.96 |
| Rotamer outliers (%) | 0.00 |
| C-beta outliers | 0 |
| Molprobity score | 0.79 |
| Ramachandran plot | |
| Favoured (%) | 98.73 |
| Outliers (%) | 0.00 |
| RMS deviations | |
| Bond lengths (Å) | 0.5 |
| Bond angles (°) | 0.5 |

**Table S2: Extended CCMV at pH 7.6 in a UV irradiated sample without photoacid data collection parameters, and reconstruction and modelling statistics.**

| | **Conventional** | **Revitrified** |
|---|---|---|
| **Data collection and processing** | | |
| Microscope | Thermo Fisher Titan Krios | Thermo Fisher Titan Krios |
| Detector | Falcon 4 | Falcon 4 |
| Energy Filter | SelectrisX, 10 eV band width | SelectrisX, 10 eV band width |
| Magnification | 165,000 | 165,000 |
| Accelerating voltage (kV) | 300 | 300 |
| Total electron exposure ($e^-/Å^2$) | 50 | 50 |
| Defocus (µm) | -0.3 – -0.9 | -0.3 – -0.9 |
| Defocus increment (µm) | 0.1 | 0.1 |
| Pixel size (Å) | 0.726 | 0.726 |
| Initial movies (no.) | 3,383 | 6,200 |
| Symmetry imposed | I | I |
| Initial particle images (no.) | 203,268 | 62,045 |
| Final particle images (no.) | 79,910 | 18,822 |
| Map resolution (Å) | 4.1 | 4.3 |
| FSC threshold | 0.143 | 0.143 |
| **Refinement** | | |
| Map sharpening B factor ($Å^2$) | 153.5 | 143.9 |

**Table S3: Extended CCMV in a sample with the photoacid released and partially contracted CCMV after revitrification, data collection parameters, and reconstruction and modelling statistics.**

| | **Conventional** | **Revitrified** |
|---|---|---|
| **Data collection and processing** | | |
| Microscope | Thermo Fisher Titan Krios | Thermo Fisher Titan Krios |
| Detector | Falcon 4 | Falcon 4 |
| Energy Filter | SelectrisX, 10 eV band width | SelectrisX, 10 eV band width |
| Magnification | 165,000 | 165,000 |
| Accelerating voltage (kV) | 300 | 300 |
| Total electron exposure ($e^-/Å^2$) | 50 | 50 |
| Defocus (μm) | -0.3 – -0.9 | -0.3 – -0.9 |
| Defocus increment (μm) | 0.1 | 0.1 |
| Pixel size (Å) | 0.726 | 0.726 |
| Initial movies (no.) | 10626 | 7902 |
| Symmetry imposed | I | I |
| Initial particle images (no.) | 139,274 | 31,521 |
| Final particle images (no.) | 30,095 | 21,675 |
| Map resolution (Å) | 3.9 | 8.0 |
| FSC threshold | 0.143 | 0.143 |
| **Refinement** | | |
| Map sharpening B factor ($Å^2$) | 133.1 | 1017.9 |
| **Model composition** | | |
| Non-hydrogen atoms | 3307 | n/a |
| Protein residues | 441 | n/a |
| Ligands | n/a | n/a |
| **Validation** | | |
| All-atom clashscore | 8.12 | n/a |
| Rotamer outliers (%) | 0.28 | n/a |
| C-beta outliers | 0.00 | n/a |
| Molprobity score | 1.66 | n/a |
| Ramachandran plot | | |
| Favoured (%) | 96.55 | n/a |
| Outliers (%) | 0.00 | n/a |
| RMS deviations | | |
| Bond lengths (Å) | 0.31 | n/a |
| Bond angles (°) | 0.51 | n/a |
